# Oncofly: A CURE for Cancer

**DOI:** 10.1101/2021.01.07.425757

**Authors:** Floyd A. Reed, H. Gert de Couet

## Abstract

Course-based Undergraduate Research Experiences (CURE’s) are emerging as a means to engage large numbers of undergraduate students in meaningful inquiry-based research activities. We describe here a simple laboratory exercise as part of an undergraduate genetics course that illustrates the contributions of oncogenes and tumor suppressors to the formation of neoplasms in an invertebrate model system. In addition, students were challenged to investigate whether flies reared on a diet containing a variety of additives display a higher number of invasive tumors in the larval abdomen.

The goal of the exercise was to (i) familiarize students with the multigenic origin of the cancer phenotype, to (ii) introduce some of the fundamental molecular cancer hallmarks, and to (iii) highlight the significance of invertebrate model systems in biomedical research. Furthermore, (iv) students learn to execute a molecular test for transgenic produce and (v) apply statistical tools to test a simple hypothesis.

We evaluated student learning and changes in opinions and attitudes relating to environmental versus genetic causes of cancer and several common misconceptions using a questionnaire before and after completing the exercise.

Overall, significant improvements in the rate of factually correct responses and reductions in uncertainty were demonstrated. Although resistance to change was apparent in regard to identifying some risk factors, there was clear learning and understanding of the core concepts of carcinogenesis and the utility of basic research with model organisms.

## Introduction

We teach that science is based upon replicable research that builds both a descriptive and predictive mechanistic model of the world around us, should embrace critical thinking, and not be dogmatic. Ideally students would be directly exposed to self guided research questions; however, in practice students are often given lists of facts to memorize and participate in “canned” laboratory exercises where the expected outcome is already known. This falls far short of teaching how science is actually done and does not set the stage to foster creativity, critical thinking, model building, and hypothesis testing (Handelsman *et al*. 2004). Inquiry-based research opportunities for undergraduates exist for a minority of students. This practice is partly driven by the objective to create an educational pipeline that leads a fraction of undergraduate students to graduate school or a medical career. The benefits of hands-on, inquiry-based research experiences for the training of a future biomedical work force are undisputed (e.g., Russell *et al.*, 2007). Limiting this to advanced special courses or participation in a research laboratory selects for a cadre of students who focus on a specific educational trajectory and a major early in their coursework and excludes the majority of undergraduates. It is recognized that most institutions lack the resources to provide these opportunities for a sizable portion of undergraduate majors (Wood 2003; Desai *et al*. 2008; Harrison *et al*. 2011, cited in Auchincloss *et al*. 2014; Brownell *et al*. 2015). Course-based Undergraduate Research Experiences (CURE’s) are designed to address this disconnect and give a broader range of students an opportunity to conduct research in the classroom (Wei and Woodin 2011; Auchincloss *et al*. 2014; Jordan *et al*. 2014; Brownell *et al*. 2015). CURE’s have been shown to have a wide range of positive effects such as increasing graduation rates and improving the interpretation of data (Goldey *et al*. 2012; Brownell *et al*. 2015; Rodenbusch *et al*. 2016).

Cancer biology is typically a specialty topic only offered to graduate and medical students because of subject complexity and relevance to therapeutic intervention and treatment. However, cancer biology can be instrumental in teaching fundamental concepts of cell-and molecular biology. Practically all aspects of cellular function—metabolism, cellular signaling, epigenetics, DNA replication and repair—are impacted by cancer. Here we argue that cancer biology can be utilized as a powerful undergraduate teaching opportunity.

In teaching biology, topics that are relevant to the students’ lives outside of academia help capture interest and build upon a preexisting conceptual framework. Essentially everyone has been impacted by cancer on some level at some point in their lives (ACS 2014) making the topic an immensely personal one. Student surveys conducted early in a genetics course (from 2011 to 2017, BIOL 375, Dept. of Biology, UH Mānoa) consistently found high interest in learning more about ‘genetically modified organisms’ and ‘cancer genetics’. In addition, there is widespread persistent misinformation about the causes and treatments of cancer (Shahab *et al*. 2018). People care about cancer and this interest can be used to effectively communicate fundamental principles of genetics and cell biology as well as teach sophisticated genetic tools, the value of model organisms and basic research, address common misconceptions and controversial issues, and even place cancer and genetics into both a social and evolutionary context.

Here we present a teaching laboratory module designed for life science undergraduates which uses a range of state-of-the-art genetic tools to generate benign tumors in the *Drosophila melanogaster* model. *Drosophila melanogaster* also simply referred to as Drosophila, has been a cancer model for over a century (Villegas 2019) and Drosophila is a tractable system in CURE settings (Chen *et al*. 2005). The student exercise described here is based on a study by Pagliarini and Xu (2003), which used Drosophila to generate tumors in the optic lobes of the fly by “two hit” over-expression of an oncogenic variant of the *Ras* gene in a background of cells deprived of the *scrib* tumor suppressor gene product by a loss-of-heterozygosity. Using the technique of mosaic analysis with a repressible cell marker (MARCM), these clones express a green fluorescent protein and tumors arising from these clones are easily visualized *in vivo*.

A critical point in development of cancer is the transition from a benign to a malignant tumor. This occurs when the tumor gains the ability to metastasize. Some of these ‘engineered’ Drosophila neoplasms seed secondary tumors in the thorax and abdomen of fly larvae—an insect analogue of metastasis. (The question of the validity of this term in relation to the satellite tumors found in the Drosophila model is discussed later.) This cellular behavior can be used to screen for substances that either promote or suppress this process (Willoughby *et al*. 2013). The effects of food additives and treatments, which can be discussed and selected collectively by the students, upon the rates of metastases are quantified in blind experiments. The data are shared with the class, statistically tested in aggregate, and reported by the students. The guided selection of treatments can follow current events regarding cancer biology. The equipment needed fits within most college laboratory budgets and can also be used in a range of other biology teaching projects. The experiment consists of a single cross and can easily be conducted by the students in a few calendar weeks. Because of the complexity of the strains used and the underlying mechanisms the exercise can easily be adapted to different educational levels and goals.

In the initial version of this experiment we combined a CURE centered on analyzing local produce for genetic modification with the introduction of principles of carcinogenesis in the expectation that it would facilitate discussions among the students about the scientific basis of claims regarding carcinogenicity of GM foods (Séralini *et al*. 2012; Casassus 2013; Grunewald and Bury 2013; Hammond *et al*. 2013; Hayes 2013; Panchin 2013; Romeis *et al*. 2013; Goldstein 2014; Wallace 2014).

Papaya is one of the major agricultural products of the State of Hawai‘i. In the mid 1990’s an epidemic of Papaya Ringspot Virus devastated the main production areas on the island of Hawaiʻi. A transgenic variant of papaya expressing a fragment of the viral capsid protein gene proved to be resistant against the disease and rescued the industry (Gonsalves 1998; Ferreria *et al.* 2002; Kallis 2013). Today, the transgenic “Rainbow” cultivar represents the majority of Hawai‘i’s papaya crop. The genome of the transgenic variant was sequenced in 2008 (Ming *et al*. 2008), which played a major role in broadening the export market beyond the North American continent by defusing concerns about the generation of immunogenic proteins as a result of inserting an open reading frame into the plant genome.

The initial GMO-focused CURE was broadened from genetically modified papaya to testing additional food additives, from health drinks to styrofoam, over the following semesters. This underscores the teaching modules’ flexibility to address a wide range of questions. We observed a surprising result; butyrate—generally considered to be protective against cancer in humans—raised the apparent rate of insect metastases. There were significant increases in students’ understanding of cancer and cancer related topics; however, students’ responses regarding risk factors and causes of cancer, although overall significantly improved according to the prevalent current scientific literature, remained resistant to change among a large fraction for topics such as GM food, power lines, and cell phone use. Students also appeared to suffer from lower retention of quantitative rather than qualitative facts, which is an area that could benefit from further exploration. Finally, the utility of model organisms to study medical phenomena such as cancer was widely appreciated with a 99.8% positive response after completing the module.

## Results and Discussion

### Experimental Results

Six food additives were double-blind tested over three semesters of teaching the module. Students and in-class instructors did not know which flies had been exposed to the treatment and which were controls until after scoring. This type of data can readily be analyzed in a 2×2 *χ*^2^ contingency table of independence—the students should have some familiarity with the *χ*^2^ test as it is often used in genetics classrooms for data analysis. No significant effect was detected on the rate of metastases with four substances: GMO versus non-GMO papaya (*Carica papaya*), commercial noni juice (*Morinda citrifolia*), multivitamin dietary supplements (“Telovites”), and polystyrene used in food packaging. Two substances resulted in a significant (using a standard *P*<0.05 cutoff) elevation of the rate of metastases: a glyphosate-based herbicide (Roundup) and butyrate (see Table 1).

**Table 1.** Summarized results of treatments and insect metastases ordered by statistical significance. Relative change in the rates of metastases are given for treatments with a *P*-value less than 0.05. All χ2 tests had one degree of freedom. The GMO papaya was compared to a control with the same amount of non-GMO papaya added. Temperature was compared by rearing Drosophila at 19 C versus 23 C. All other tests were done without the food additive added to the control food. * indicates results that are significant with a p<0.05. ** indicates results that are significant after a multiple testing correction over all experiments.

| Additive | Relative change | Sample Size | $\chi^2$ value | $P$ value |
| --- | --- | --- | --- | --- |
| Noni juice | - | 322 | 1.49 | 0.22 |
| Multivitamin | - | 275 | 1.68 | 0.19 |
| GMO Papaya | - | 481 | 2.94 | 0.086 |
| Polystyrene | - | 372 | 3.406 | 0.065 |
| Roundup | 4.9% | 360 | 3.99 | 0.046 * |
| Butyrate | 10.5% | 418 | 13.2 | $2.8 \times 10^{-4}$ ** |
| Temperature | 28.3% | 1168 | 124.75 | $1.2 \times 10^{-6}$ ** |

There was also a significant effect of temperature. The GAL4/UAS system used in this cross is temperature sensitive (Duffy 2002). When crosses are conducted at 23 C versus 19 C there is a significant elevation of mestatases. This underscores the need to run a randomly placed control in parallel during the course of each new experiment in order to account for environmental fluctuations.

It is also appropriate to point out to the students—especially when presenting the results from past semesters—the issues of multiple testing and replication. It is expected that by chance one out of twenty tests will have a *P*-value of less than 0.05 if the test used is appropriate and there is no actual mechanistic effect of the treatment (a type I error). A Bonferroni correction (e.g., Cabin and Mitchell 2000), while conservative, is the easiest to initially understand and use. It keeps the total false positive rate at 5% by multiplying each *P*-value by the total number of tests done. In this example only butyrate and temperature have significant effects after the correction. Finally, none of these experiments have been replicated outside of a single semester classroom. Replication of an earlier experiment, especially given the potential sensitivity to environmental conditions, and subsequent meta-analysis should also be proposed to the students as an important choice for testing.

Many of the substances tested fall squarely within developing results and debate—the reason for selection—which we argue enhances students’ interest in analyzing and communicating the results. The carcinogenic potential of glyphosate is of particular current focus because of the controversy surrounding recent court cases (e.g., Charles 2019, Peterson 2019) and the results found by the students illustrate the degrees of uncertainty inherent in scientific research, which can increase interest in science (Retzbach and Maier 2015). Claims about the health effects of noni (*Morinda citrifolia*) have also been debated (e.g., Brown 2012; Schulz 2014) and there are persistent statements about concerns regarding GMO food and cancer (e.g., Barrell 2019) as well as plastics, including styrene and polystyrene/styrofoam (e.g., Fox 2019; Christensen *et al*. 2018; with broader concerns about food containers that can be tested in the future, e.g., Soto and Sonnenschein 2010; Liao and Kannan 2013). The multivitamin was chosen to be tested because it claims on its website that “TeloVite, [is] the first telomere lengthening multivitamin” and that it contains antioxidants (https://www.westmartinlongevity.com/products/telovite/). Concerns can be found about possible negative effects of multivitamins and food supplements in general (e.g., Anonymous 2018; Chen *et al*. 2019), antioxidants and metastasis (e.g., Piskounova *et al*. 2015), and the relationship between telomere length and replicative potential of cancer cells is well documented (e.g., Blasco 2005; Williet *et al*. 2010). Butyrate is thought to be protective against the formation of some human cancers and combines current interests in understanding both microbiotic and epigenetic effects (Donohoe *et al*. 2012; Bultman 2017).

### Student Results

Students were given identical questionnaires before starting the “oncofly” module and after its completion (Supplementary material 1). Students were instructed to give their candid individual response to the questions, omit their names or identifying information, and that they recieved full credit for turning in a completed form regardless of responses. We were also interested in the effects of the laboratory project on their general knowledge of cancer rather than teaching them how to answer a limited set of specific questions. We were intentionally passive in presenting this information. The answers are contained in a handout (Supplementary material 2) and assigned reading materials, and aspects are touched upon both in lab presentations and an associated classroom lecture. However, students were not prompted to memorize the answers to these specific questions.

One part of the questionnaire asked students to rate statements on a scale from false (1) to uncertain (3) to true (5). The responses are summarized in Tables 2 and 3.

**Table 2.** The keyword is only used here to guide comparisons between tables 2 and 3. It did not appear on the student questionnaire.

| Keyword | Question |
| --- | --- |
| Sharks | Sharks and insects do not get cancer. |
| Artificial | Cancer only results from exposure to artificial substances. |
| Organic | Cancer can be avoided by only eating organic food. |
| Environment | Cancer only results from exposure to toxins or radiation in the environment. |
| GMOs | GMO food is known to cause cancer. |
| Cell phones | Cell phones cause cancer. |
| Power lines | Power lines cause cancer. |

**Table 3.** Statistical summary of the student questionnaire. Pre and Post refer to completing the questionnaire before and after the module. n denotes number of students. The sample sizes can vary because of differences in student attendance, and some responses were incomplete. The percent with correct answers are given first and the proportion of uncertain responses are given next. 1. *P*-value from a Wilcoxon rank sum test. 2. *P*-value from a two-proportion z-test.

| Keyword | Pre-n | Post-n | Pre-<br>correct | Post-<br>correct | Correct P-value (1) | Pre-<br>uncertain | Post-<br>uncertain | Uncertain<br>P-value (2) |
| --- | --- | --- | --- | --- | --- | --- | --- | --- |
| Sharks | 578 | 575 | 61.2% | 78.6% | $2.2 \times 10^{-16}$ | 20.2% | 7.30% | $1.9 \times 10^{-10}$ |
| Artificial | 422 | 420 | 93.4% | 97.1% | $2.0 \times 10^{-7}$ | 4.50% | 1.90% | 0.032 |
| Organic | 578 | 576 | 91.2% | 91.7% | 0.033 | 5.36% | 5.38% | 0.989 |
| Environment | 417 | 420 | 89.2% | 89.5% | $5.3 \times 10^{-3}$ | 5.04% | 4.29% | 0.607 |
| GMOs | 578 | 576 | 52.1% | 64.6% | $2.2 \times 10^{-5}$ | 33.6% | 24.1% | $4.1 \times 10^{-4}$ |
| Cell phones | 595 | 590 | 35.0% | 58.0% | $2.2 \times 10^{-16}$ | 31.8% | 24.4% | $4.8 \times 10^{-3}$ |
| Power lines | 591 | 594 | 45.7% | 64.6% | $6.7 \times 10^{-16}$ | 39.4% | 26.1% | $1.0 \times 10^{-6}$ |

**Table 4.** The keyword is only used here to guide comparisons between tables. It did not appear on the student questionnaire.

| Keyword | Question |
| --- | --- |
| Carcinogen | Can you name a known carcinogen that we are regularly exposed to? |
| Gene types | What are the two types (categories) of genes that play a role in cancer? |
| Cause | Do you know the leading cause of cancer in the US? |
| Gene name | Can you name a cancer gene? |
| Natural | Do you know any natural carcinogens? |

**Table 5.** Uncertain answers (no or equivalent) were included with incorrect answers in quantifying and testing the proportion correct with a two-proportion z-test. The proportion of uncertain responses out of the total were also tested with a two-proportion z-test.

| Keyword | Pre-n | Post-n | Pre-<br>correct | Post-<br>correct | Correct P-value | Pre-<br>uncertain | Post-<br>uncertain | Uncertain<br>P-value |
| --- | --- | --- | --- | --- | --- | --- | --- | --- |
| Carcinogen | 600 | 598 | 42.3% | 54.2% | $4.1 \times 10^{-5}$ | 33.8% | 21.7% | $3.0 \times 10^{-6}$ |
| Gene types | 601 | 600 | 4.66% | 49.5% | $2.2 \times 10^{-16}$ | 65.2% | 14.2% | $2.2 \times 10^{-16}$ |
| Cause | 600 | 600 | 37.7% | 65.5% | $2.2 \times 10^{-16}$ | 28.3% | 10.7% | $1.1 \times 10^{-14}$ |
| Gene name | 600 | 597 | 20.7% | 64.2% | $2.2 \times 10^{-16}$ | 63.5% | 12.1% | $2.2 \times 10^{-16}$ |
| Natural | 600 | 601 | 41.0% | 56.7% | $4.9 \times 10^{-8}$ | 47.7% | 26.8% | $7.2 \times 10^{-14}$ |

**Table 6.** The keyword is only used here to guide comparisons between tables. It did not appear on the student questionnaire.

| Keyword | Question |
| --- | --- |
| 5 Year | What is the five year survival rate for all cancers combined in the US? |
| Lifetime | What is a person's chance of being diagnosed with cancer at some point in their lives? |
| Hereditary | What fraction of cancer cases are familial or hereditary (passed down in families due to genetic factors)? |

**Table 7.**
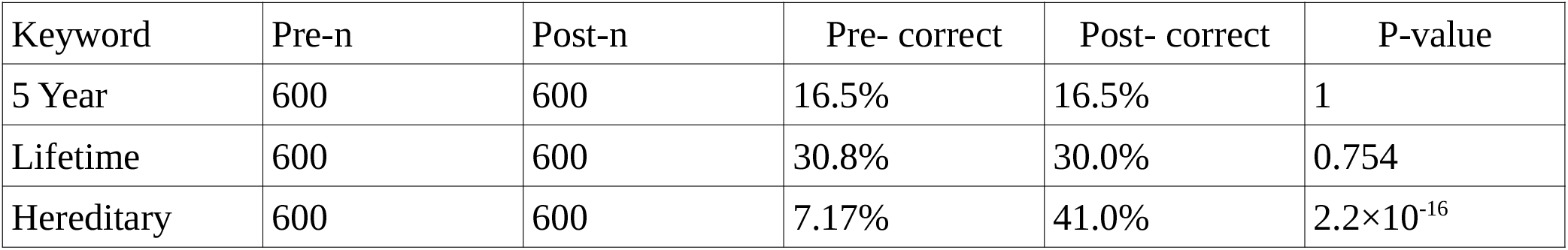
The change in proportion of correct responses out of the total was tested with a two-proportion z-test.

| Keyword | Pre-n | Post-n | Pre- correct | Post- correct | P-value |
| --- | --- | --- | --- | --- | --- |
| 5 Year | 600 | 600 | 16.5% | 16.5% | 1 |
| Lifetime | 600 | 600 | 30.8% | 30.0% | 0.754 |
| Hereditary | 600 | 600 | 7.17% | 41.0% | $2.2 \times 10^{-16}$ |

The rate of correct, according to the current prevalent scientific literature, volunteered answers significantly improved for every question asked after participating in the oncofly project. Uncertainty of the correct answer declined significantly for the majority of questions asked and in no case increased significantly. For example, the proportion of students disagreeing with “GMO food is known to cause cancer” increased from 52.1% to 64.6% and uncertainty regarding the answer to this question decreased from 33.6% to 24.1%. A summary of these results are plotted in figure 1.

**Figure 1.**
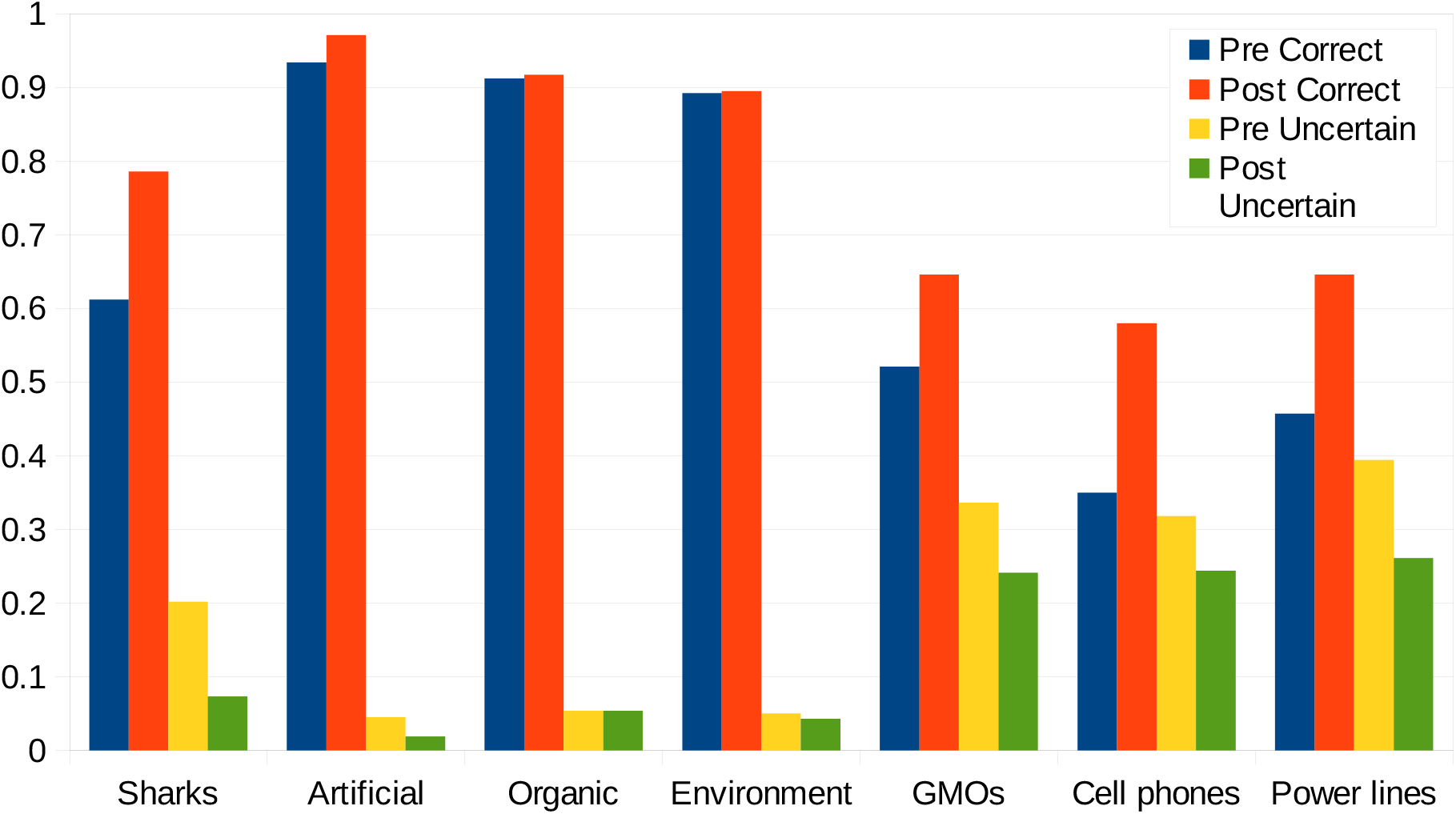
Plot of the results of correct and uncertain responses before and after the experiment. Data is detailed in Table 1.

**Figure 2.**
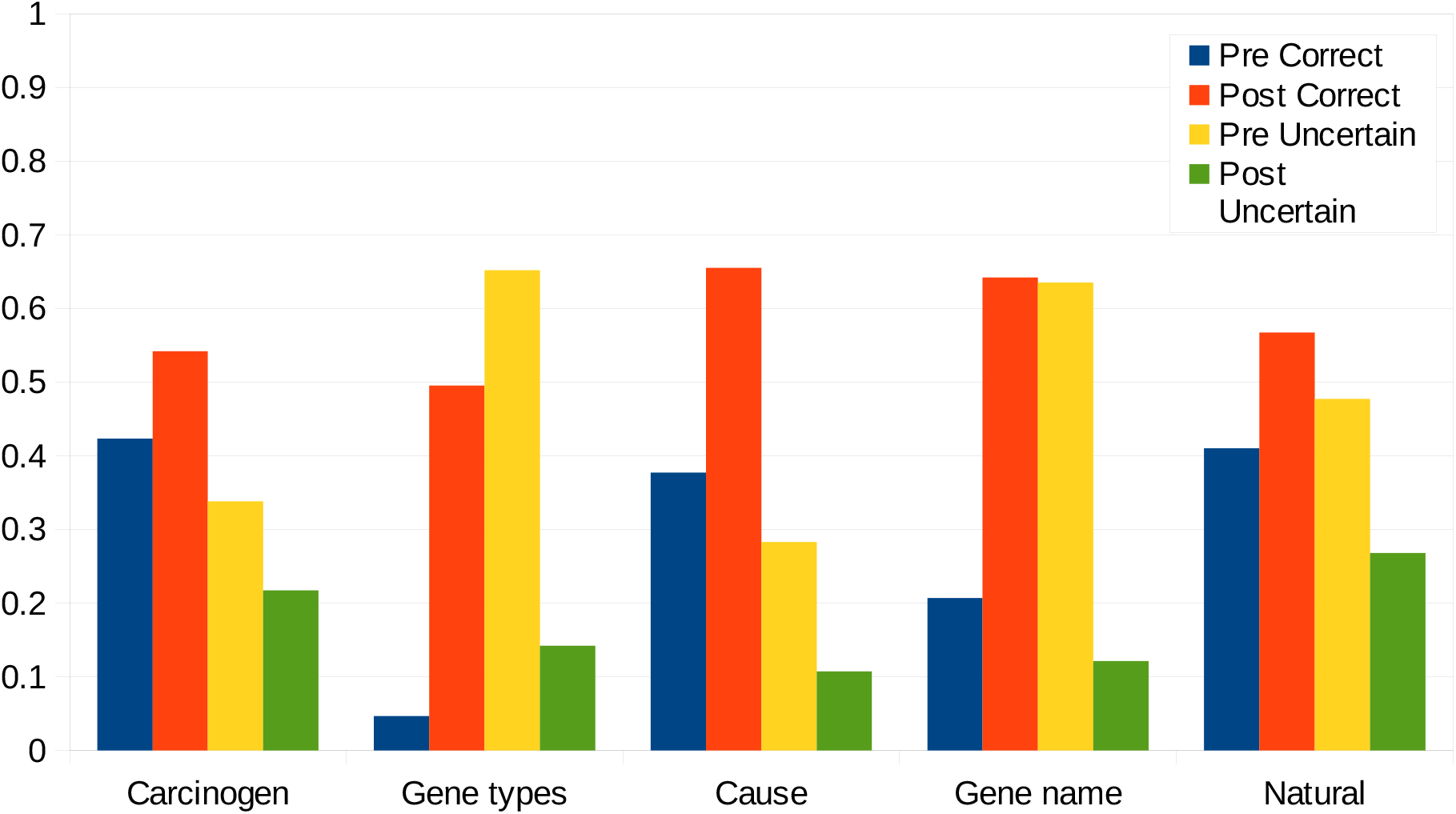
Plot of the results of correct and uncertain responses before and after the experiment. Data is detailed in Table 4.

Students were also asked a series of short answer questions to test recalling specific information.

In summary, there was a significant reduction in uncertainty and increase in correct cancer-relevant information recall for all of these five questions.

Correct answers to naming a carcinogen to which we are regularly exposed increased from 42% pre to 54% post. This may seem frustratingly low, but many of the incorrect answers were substances such as asbestos, which are carcinogens but not ones to which we are regularly exposed. The most common single incorrect answer was nicotine, which is not conclusively a carcinogen (Haussmann and Fariss 2016 see also Chernyavsky *et al*. 2015). This is a common misconception (e.g., Wilson *et al.* 2011) that we plan to address in future classes. The most common correct answers were UV exposure from sunlight (or equivalent, e.g., the sun) and tobacco smoke (or equivalent, e.g., cigarettes).

Similarly, the correct response of naming the two categories of cancer genes, (proto)oncogenes and tumor suppressors, increased from 4.6% pre to 50% post (pre-n 601, post-n 600, *P* = 2.2×10^−16^,). Many of the answers in the post survey were close but frustratingly incorrect. Examples included only giving one or the other category correctly, e.g., proto-oncogene and oncogene, being overly gene specific e.g., *Ras* and *scrib*, or category specific e.g., cell growth and DNA repair.

The correct response for the leading cause of cancer, tobacco smoke or equivalent increased dramatically from 38% to 66%. Lung cancer and nicotine were not counted as correct for this question. Also, correct answers for naming a gene involved in cancer increased from 21% to 64%. *BRCA1* and *Ras* were common responses followed by *p53* and *scribbled*. Oncogene, tumor suppressor, as well as only “BRCA”, were common incorrect answers. Perhaps this question should be changed to “can you name a specific cancer gene” in the future to reduce ambiguity.

Correct answers for natural carcinogens increased from 41% to 57% (pre-n 600, post-n 601, P = 4.9×10^−8^, two-proportion z-test). Common correct responses were UV from sunlight, tobacco, aflatoxin, and asbestos. Common incorrect answers were nicotine, cigarettes, carbon monoxide, and methane. This is a question with some surprising misconceptions (carbon monoxide and methane) and highlights and area that can be expanded upon in providing the students with additional information.

In contrast to the previous responses, quantitative proportions appeared to be more difficult for the students to extract and/or retain from the lecture and reading.

The one question out of three with a significant improvement focused on genetic factors. Since this was conducted as a part of a genetics class the students may have been primed to retain genetic information as a byproduct of the other topics discussed in class. While not the goal of this work, and this is a small sample of questions, it raises an interesting contrast and asks how this can be addressed in the classroom. The challenges of developing quantitative skills among biology majors has been widely recognized and ways to teach and enhance quantitative skills among biology students is an active area of inquiry (e.g., Speth *et al*. 2010; Gormally *et al*. 2012; Hester *et al*. 2014). Perhaps this extends beyond performance on a specific task to more passive use of quantitative information from lectures and reading.

We also asked yes or no if “the following is a risk factor for cancer”.

Only two had a significant change, cellphone use and obesity, both with an increase in correct answers based on current scientific knowledge (compare cellphone use as a risk factor, Table 8, to cellphones cause cancer, Table 3; anecdotally there seemed to be an over-generalization among students that “radiation” caused cancer rather than ionizing radiation and UV light). Two factors were already predominantly correct in the pre-survey and thus had less power to detect a change. The glaring exception is chronic inflammation, which is a key component of cancer development (Coussens and Werb 2002; Hanahan and Weinberg 2011) and presents an additional point to focus on presenting to the students.

**Table 8.** The change in proportion of positive responses out of the total was tested with a two-proportion z-test.

| Factor | Pre-n | Post-n | Pre- “yes” | Post- “yes” | P-value |
| --- | --- | --- | --- | --- | --- |
| Age | 600 | 600 | 87.2% | 89.0% | 0.327 |
| Cellphone use | 600 | 598 | 49.3% | 35.2% | $6.78 \times 10^{-7}$ |
| Chronic inflammation | 600 | 600 | 44.7% | 49.2% | 0.118 |
| Obesity | 601 | 599 | 56.8% | 65.8% | $1.37 \times 10^{-3}$ |
| Power lines | 600 | 600 | 21.8% | 19.8% | 0.394 |

Again, we presented information specific to these questions in an indirect manner to broadly test for improvement in knowledge about cancer and cancer genetics. Instructors may wish to provide and test this information directly and inform students that they must know the answers to these specific questions between pre- and post- assessment. This will likely result in greater improvements for these specific questions; however, we feel that this does not do as well at assessing broader knowledge outside of the assigned questions. Ultimately the best strategy may be a mixed approach to assess both specifically assigned information and broader knowledge.

Finally, a question with a completely open ended component was asked “Are model organisms (like mice or Drosophila fruit flies) useful tools for studying cancer? Why or why not?” The proportion of positive responses increased from 89.9% pre- to 99.8% post- (pre-n 584, post-n 597, χ^2^ = 60.4, d.f.= 1, P = 7.6×10^−15^). Among the reasons given for negative responses were short life span of model organisms, genetic differences between model organisms and humans, the same risk factors cannot be studied, human specific cancer cannot be studied, flies cannot get cancer, carcinogens vary between species, and it is unethical to induce cancer in other organisms. Among the reasons given for positive responses were genetic and cellular similarity between humans and model organisms, model organisms are easy and cheap to maintain, short life cycles, sequenced genomes, ease of genetic manipulation, it is more ethical to study cancer in model organisms than studying it in humans, the ability to rear large numbers of genetically uniform individuals for testing and avoiding the problems of genetic heterogeneity and incomplete penetrance in human cancer cases.

Model organisms tend to be small and short lived. In many ways cancer is a disease of age, accumulated DNA damage from the environment, and possibly body size. Indeed, recent insights have come from studying a range of non-traditional long-lived species (see Supplemental Discussion). Some students become aware during the course of the experiment of the discrepancy that organisms with short lifespans and limited cell divisions are used to study cancer. This is the reason causative genes in the fly are targeted in order to achieve neoplastic transformation within a short lifespan. This is not to detract from model organisms (the Drosophila model has resulted in a wide range of discoveries in cancer research, Sonoshita and Cagan 2017), but the unique circumstances of each species must be kept in mind. The critical transition to a metastasizing malignant tumor in humans is thought to require additional mutations and is characterized by typical changes in gene expression, specifically the upregulation of genes that are involved in cell motility and downregulation of cell adhesion molecules (e.g., Barrallo-Gimeno and Nieto 2005; Birchmeier 2005; Sedgwick and D’Souza-Schorey 2016). It is not clear that these changes occur in tumors of the fly abdomen in this experiment; however, an ambitious class might try to test this via quantifying relative RNA abundance of candidate genes. The butyrate treatment result of an increase in abdominal tumors may suggest an alternative. Butyrate is a histone deacetylase inhibitor. In humans this is thought to help prevent certain forms of cancer (Donohoe *et al*. 2012). The initial benign tumors in this experiment are limited to the head region because of their dependence on *eyeless* expression of the *flippase* recombinase to result in mitotic recombination and loss of heterozygosity of the tumor suppressor *scribbled*. Histone modifications are a mechanism of determining cell fate. Disruptions to epigenetic programming can both disrupt normal gene expression at a subset of genes and directly contribute to a cancer phenotype (e.g., Hirabayashi and Gotoh 2010; Jones *et al*. 2016). These abdominal tumors may not be satellite tumors arising from metastasis of a tumor in the head but rather benign tumors arising *in situ* from the activation of *flippase* via epigenetic dysregulation. If this is the case, which remains to be seen, the fly model could prove useful in screening substances that promote tumor formation through epigenetic means (Momi *et al*. 2014). Regardless of the degree of similarity between the Drosophila model and human cancer, this module does serve to expose students to the most important fundamental components of cancer genetics and coveys how to use genetics tools to study these processes.

This work was originally inspired by the GMO debate and we include methods for testing the GMO status of plants by PCR. Suggestions for presentation and discussion of the relationship or lack thereof of cancer and GMOs are provided in the Supplemental Discussion.

Another common misconception apparent among the students was the belief of a link between some forms of non-ionizing electromagnetic radiation (cell phones, power lines) and cancer. This is an area of active research and controversy (Hertsgaard and Dowie 2018; Landler and Keays 2018; NCI 2019). In the spirit of this CURE, there are a range of creative ways to for classrooms to test electromagnetic radiation using the Drosophila cancer model.

In addition to illustrating the utility of model organisms, in this case *Drosophila melanogaster*, cancer biology underscores the importance of pure research. This can be summarized by three Nobel prizes in Physiology or Medicine. Peyton Rous was interested in natural history and later pathology. He investigated a tumor in a hen brought by a person to the institute where he worked. He could not have known that his work would identify oncogenic viruses and lead him on a path that also identified chemical carcinogens, two distinct mechanisms of carcinogenesis, and ultimately resulted in the 1966 Nobel Prize (Weiss and Vogt 2011). Tim Hunt was working on getting sea urchin eggs to divide without being fertilized and noticed an odd pattern of a protein that appeared and disappeared with each cell division. In light of the current difficulty in obtaining funding for basic research it is sobering to read Hunt (2015) and Robertson (2015), also listen to Al-Khalili (2011) for an audio interview. There were a series of chance events that led to the discovery of cyclins that control the cell cycle, difficulty in getting the scientific community to realize the significance, had implications for understanding cancer, and ultimately resulted in the 2001 Nobel Prize. More recent breakthroughs have been in the interface between the immune system and cancer, which resulted in the 2018 Nobel Prize. ‘Neither of the two Nobel Prize winners, Jim Allison and Tasuku Honjo, directly set out to cure cancer – “that wasn’t it at all,” Allison has said – they were trying to understand how the immune system works’ (Davis 2020). In the process the total rates of cancer in the US peaked in 1991 and have since fallen (13% from 1991 to 2015, NCI 2018) deaths due to cancer have also fallen (29% from 1991 to 2017, Siegel *et al*. 2020), and five year survivorship has increased (26% from 1991 to 2015, NCI 2018). A number of new tools are continuing to be developed. We should recognize the breakthroughs that stem from fundamental biological research and that none of these leaps in understanding would have happened if there were not people trained in biological research. In order to understand and address cancer, support biology research and education.

## Acknowledgements

We thank the students and teaching assistants of BIOL 375L Genetics Lab, University of Hawai‘i at Mānoa for participating in this project. We thank Sachie Etherington and Natasha Isaac for coordinating the teaching lab. We thank Zhaotong Xu, Tahoora Pourjalali, Herena Ha, and Keisha Willis for help maintaining the fly stocks and media preparation. We thank Vanessa Reed for editing and proofreading. David Bilder, Ryan Boileau, Tian Xu, and La-Di Ming graciously provided Drosophila stocks.

## Methods

### Research regulation

This is written from the perspective of regulation within the United States; check for equivalent regulations in your own jurisdictions. Research in a classroom setting fall into a regulatory gray zone (e.g., Tomkowiak and Gunderson 2004; Callier 2012). This is also the case for using transgenic organisms in a classroom. NIH guidelines do not mention classroom settings and education in their transgenic biosafety rules (https://osp.od.nih.gov/biotechnology/nih-guidelines/). Individual institutions, as the University of Hawai‘i administration has done, may choose to regulate this teaching module as laboratory research. In this case this work is registered under IBC protocol no. 18-08-932-04-1A. You should check with your institute’s biosafety committee (EHSO, Environmental Health and Safety Office or equivalent) for guidance regarding classification of lab activities with genetically modified organisms.

### Fly stocks and rearing

A CO_2_ based fly anesthetization system (a pressurized CO_2_ source, needle, pad, paintbrushes, dissecting microscope, *etc*.) is highly recommended for working with Drosophila in the classroom. Fly stocks are maintained on standard yeast-glucose agar media in trays of vials with cotton stoppers.

Two complex genetic lines of flies, designed to model tumor metastasis, are crossed together:

Line 1: w[-]; UAS-Ras[V12]; FRT2A, FRT82B Scrib[-]/TM6B

Line 2: y[-], w[-], ey-FLP; Act-Gal4; UAS-GFP S56T; FGT82B; Tub6-Gal80

These lines make use of the GAL4/GAL80/UAS expression control system (Duffy 2002) and the FLP/FRT system (Golic and Lindquist 1989; Wu and Luo 2006) activated in the larval head to make double stranded DNA cuts at specific chromosomal positions resulting in mitotic recombination. Furthermore, the oncogene Ras[V12] gain-of-function hypermorph is expressed simultaneously with a loss of heterozygosity, as a result of mitotic recombination, of a tumor suppressor scrib[-] amorph which triggers precancerous and cancerous cell behavior. The loss of heterozygosity also results in the expression of Green Fluorescent Protein (GFP) which marks the precancerous cells. GFP expressing cells were visualized with a Nightsea stereo-microscope fluorescence adapter (https://www.nightsea.com) with a 440–460 nm excitation light and 500–560 nm emission filter. A MiniVID eyepiece camera was used to record examples.

See Supplemental methods for the fly food recipe used.

The vials were labeled with 10 different numbers. Five numbers corresponded to controls and five numbers corresponded to the food additive to be tested. The numbers overlapped in sequence (e.g., control: 3, 4, 6, 8, 9; test: 1, 2, 5, 7, 10) The key for decoding the food was kept by the lab manager and not made available to either the students scoring the larvae or to the teaching assistants and class instructor until after the experiment had finished and the final numbers were tabulated (i.e., a double-blind experiment).

Line 1 and line 2 were crossed to each other using standard Drosophila lab technique (e.g., Greenspan 2004) and allowed to lay eggs on the food for three days before the adults were cleared from the vial. After one week the third instar larvae were scored. The food can autofluoresce so it is recommended that the larvae be washed briefly in water before scoring. The presence of a clear point or points of GFP expression in the posterior 2/3 of the larvae were recorded as a metastasis.

It is necessary that a control be run in parallel with each treatment to be tested rather than comparing to previous control results or current and prior controls in aggregate. The GAL4 system is sensitive to environmental conditions such as temperature which may vary between semesters. The use of insect incubators, which can be temperature and light controlled (e.g., 14:10 hr light/dark at 24 C), are recommended if possible.

### GMO detection

Students were asked to bring papaya samples (commonly grown here in Hawai‘i) from a wide range of sources (grocery store, farmer’s markets, homegrown) and to attempt to include plants not thought to be genetically modified. The class instructors were also able to obtain GMO and non-GMO papaya. Fruits were labeled with tape and a marker and stored in a refrigerator. A small section of the fruit skin, less than a square centimeter, was removed and DNA was extracted (Qiagen DNeasy plant kit). This was used for PCR (see Merritt *et al*. 2008 for a way to conduct PCR if a thermocycler is not available). The following PCR primers have been found to perform well in the classroom and can be ordered from online DNA oligo synthesis services (we used https://www.idtdna.com/).

Standard rbcL DNA barcoding primers were used as a positive PCR control (Kress and Erickson 2007).

rbcL-a-f 5’-ATGTCACCACAAACAGAGACTAAAGC

rbcL-a-r 5’-GTAAAATCAAGTCCACCRCG

The 35S promoter from the cauliflower mosaic virus (CaMV) is present in a wide range of genetically modified crop plants (including papaya) and provides a useful way to screen for GMO status by PCR (e.g., Hurst *et al*. 1999 and references therein).

CaMV35Sf 5’-GCTCCTACAAATGCCATCA

CaMV35Sr 5’-GATAGTGGGATTGTGCGTCA

The temperature cycling scheme 1 was used for both PCR reactions (this allows them to be run simultaneously in the same thermocycler). This temperature cycling is optimized for the 35S primer pair. If two different temperature cycling schemes are used the scheme 2 should be used for the rbcL primer pair.

Temperature cycling 1: 95 C 3 min, 33 cycles of (94 C 20 s, 54 C 40 s, 72 C 1 min), 72 C 3 min

Temperature cycling 2: 95 C 3 min, 33 cycles of (94 C 30 s, 55 C 30 s, 72 C 1 min), 72 C 10 min

The PCR reactions are designed for a 20 μl final volume. A 5X master mix is provided to the students who must dilute it with purified water and add the correct primer pair and a 1 μl DNA sample to a final 1X concentration of 200 μM dNTP mix, 1.5 mM MgCl2, and 0.05 U/μl Taq polymerase, and 1 μM primer mix.

Three PCR reactions were set up for each sample: a positive control with the rbcL primers, a negative control with 1 μl purified water substituted for the DNA sample, and the GMO-test PCR with 35S primers.

### Food additives

#### GMO and non-GMO food

Commercially available Rainbow Papaya that was confirmed by PCR to be genetically modified and a papaya from Kumu Farms, Maui that was confirmed by PCR to not be genetically modified were used. The whole fruit was washed with filtered purified water and wiped down with 70% ethanol to sterilize the surface. Then the fruits with the skin and without the seeds were pureed, approximately ½ kilogram of fruit puree was added to the Drosophila food.

#### Noni juice

375 ml of commercially available “Puna Noni Naturals Noni Juice” Noni Connection, Inc. (“100% Pure Hawaiian Noni Juice”) was used.

#### Vitamin supplement

0.08 g of ground Telovites (Telovites, West Martin, Inc.) was added. The ingredients list of the Telovites used is copied here. “Vitamin A (as acetate, beta-carotene) Vitamin C (as ascorbic acid, calcium ascorbate) Vitamin D3 (as cholecalciferol) Vitamin E (as d-alpha tocopheryl succinate and mixed tocopherols) Vitamin K2 (as MK7) Vitamin B1 (as thiamin HCl) Vitamin B2 (riboflavin) Niacin (as Niacinamide) Vitamin B6 (as pyridoxine HCl) Folic acid Vitamin B12 (as cyanocobalamin) Biotin Pantothenic Acid (as d-calcium pantothenate) Calcium (as carbonate, dibasic calcium phosphate, citrate, ascorbate) Phosphorus (as dibasic calcium phosphate) Iodine (potassium iodide, kelp) Magnesium (as oxide, citrate) Zinc (as amino acid chelate) Selenium (as selenomethionine) Copper (as amino acid chelate) Manganese (as amino acid chelate) Chromium (as polynicotinate) Molybdenum (as sodium molybdate) Chloride (as potassium chloride) Potassium (as potassium chloride) Green Tea leaf extract Standardized to 90% polyphenols Chlorella (Chlorella vulgaris) Resveratrol (providing 30 mg trans-resveratrol) Choline bitartrate Astragalus membranaceus Providing Polysaccharides 40 mg L-Carnosine Inositol Grape seed extract (85-95% OPC) Lycopene Lutein Silicon (as sodium metasilicate) Boron Vanadium (as vanadyl sulfate) Nickel (as nickel sulfate).”

#### Styrofoam

17.78 g of ground up commercially available styrofoam/polystyrene plates were added to the food.

#### Butyrate

750 mg of sodium butyrate (Sigma-Aldrich B5887-250MG) were added.

#### Roundup

12.5 ml of commercially available “Roundup Ready-To-Use Weed & Grass Killer III” (2.0% glyphosate isopropylamine salt, 2.0% pelargonic acid & related fatty acids, 96.0% other ingredients) was added to the food.

## Supplemental Discussion

### Lecture presentation suggestions

Cancer genetics is a natural extension of topics such as mitosis, cell cycle control, and gene product interactions (e.g., pRB, E2F, Cyclin D1, and Cdk4; Tyson *et al*. 2001; Davidich and Bornholdt 2008; Figure 1 of Akin *et al*. 2014) and can be introduced fairly early in a typical genetics class. There are also aspects of developmental genetics that are relevant background for cancer genetics. Cells in multicellular organisms depend extensively on signals from other cells for growth, cell fate, and cell death signals (*Notch* and *Wnt* are good examples; e.g., Artavanis-Tsakonas *et a*l. 1999; Loh *et al.* 2016). Understanding the role of the cellular environment and components like p53 and TOR in growth signals, growth inhibition, and programmed cell death is an important aspect of cancer development (Ashcroft and Vousden 1999; Fingar and Blenis 2004; Hanahan and Weinberg 2009; Hanahan and Weinberg 2011; Pacheco *et al*. 2014). Finally, cancer evasion of the immune system and new cancer therapies based on utilizing the immune system should be illustrated (e.g., Couzin-Frankel 2013; Muenst *et al*. 2016).

Epidemiological data provide insights into cancer causes and development (Ames *et al*. 1995). The change in rates of deaths due to cancer over the last century illustrate the effects of smoking on lung cancer and likely effects of changes in food preservation on stomach cancer (Peto *et al*. 2000; Figure 8 of Siegel *et al*. 2016). The 20 to 30 year lag between changes in the rates of smoking and lung cancer (NCI 2003) is especially informative as to the time it takes for cancer to develop after exposure to risk factors. This illustrates that cancer development is a long process of trial and error by mutations, not the least of which is maintaining evasion of the immune system by a subset of successful cells. Finally, cancer incidence plotted by age illustrates that cancer is also a disease of aging with rates increasing dramatically after 55 years of age (CRUK 2019).

This teaching module was originally inspired by a genetics classroom (BIOL 375) discussion of the controversial results of Seralini *et al*. (2012), which claimed that feed containing genetically modified (GM) crop plants caused cancer. This publication resulted in widespread media impact (Romeis *et al*. 2013 and references therein) and controversy (e.g., Casassus 2013). The class discussion included pointing out the disconnect between statements made in the publication and the results provided, additional information such as the rates of tumors that occur in the type of rats used, as well as psychological and social issues such as confirmation bias and framing effects in the media. Irrational fear of new technologies (e.g., cell phones, nanotechnology, artificial intelligence) among a subset of the population makes people receptive to speculation that some of these new technologies are responsible for problems with cryptic origins like cancer. While there was no mechanistic reason to infer that consumption of GM produce caused cancer (in fact quite the opposite in some cases, e.g. Munkvold *et al*. 1999; Dowd 2000), the students remained divided on the core question; with very limited sample sizes was there a reasonable amount of evidence in either direction, that GM organisms are associated with or not associated with increased cancer rates? The point was brought up that, as scientists, we should conduct a similar study and test this for ourselves in the teaching lab with more statistical power from a larger sample size.

If you and/or your students choose to test GMOs, we encourage you to also include a discussion of mechanistic reasons why specific GMO food could be harmful (e.g., Nordlee *et al*. 1996), protective (e.g., Dowd 2000; Singh and Bhalla 2008; Jean-Yves *et al*. 2017), or neither (e.g., Snell *et al*. 2012) as well as larger questions such as the idea of what is “natural” in the context of transgenics and engineered food (e.g., Dubcovsky and Dvorak 2007; Pace *et al*. 2008; Hehemann *et al.* 2010; Kyndt *et al.* 2015; Soucy *et al*. 2015) and some of the social issues involved (e.g., McGray 2002; Enserink 2008; Stone 2010; Lynas 2013; Fischer *et al*. 2015). This also has implications for the GMO labeling debate (e.g., Roff 2009; Prentice 2018), which is also an appropriate point to discuss in the classroom.

Part of the philosophy behind designing this lab was to illustrate the value of model organisms. However, recent insights have also come from studying a range of non-model species. This provides an opportunity to broaden the discussion beyond medical applications to an evolutionary context (both within tumors, Vitale *et al*. 2019, boundaries to gene flow and speciation, Schartl 2008, and across the tree of life, Aktipis *et al*. 2015; Albuquerque *et al*. 2018). First of all is the observation, known as Peto’s paradox, that large and/or long lived organisms do not have higher rates of cancer despite having more cells and opportunities for DNA damage. These organisms have evolved strategies of cancer suppression such as increased copies of p53 in elephants (Callaway 2015 and references therein), a unique extracellular environment in naked mole rats (Toole 2004; Tian *et al*. 2013), and cell walls in plants (Doonan and Sablowski 2010).

Another excellent case of cancer and evolution is found among transmissible cancers. Transmissible cancer has conservation implications in the Tazmanian Devil and helps to illustrate the dangers of low genetic diversity of the immune system (Bostanci 2005; O’Neill 2010; Woods *et al*. 2015; Caldwell and Siddle 2017). Transmissible cancer has also been found among shellfish and may be more common than realized in marine systems (Metzger *et al*. 2015; Metzger *et al*. 2016). Some transmissible cancers may be tens of thousands of years old, evolving along with their host and becoming established in new species (Murchison *et al*. 2015; Strakova and Murchison 2015). Carried to an extreme, there is a hypothesis that some parasitic animals, Myxosporea, may have evolved from transmissable cancer (Panchin *et al*. 2019).

Cancer is a breakdown of cellular cooperation in multicellular organisms (Aktipis *et al*. 2015). We can further broaden our focus and ask what cancer-like dynamics would look like in super-organisms or even social organizations. A mutant lineage in an ant species, *Pristomyrmex punctatus*, only produces queens which migrates to new colonies to live off of the workers and avoid local extinction within the host colony (Dobata *et al*. 2011); this is essentially cancer on a super-organism level with analogs of gain of function mutations, loss of regulation, and metastasis. In human societies policing is used to suppress competition and enforce cooperation in groups of individuals (e.g. Frank 2003; West *et al*. 2007); which has parallels with the roles of tumor suppressors ensuring cellular cooperation—these perspectives and analogues may aide students development of cancer concepts and inspire new directions.

## Supplemental Methods

Two batches of Drosophila food are made simultaneously, one as a control and one with a food additive to be tested, according to the following recipe.

Ingredients:

Starting water 3,125 mL

Hydrating water 625 mL

(Organic, non-GMO) Corn flour 272.5g

Yeast 47.5g

Sucrose 80g

Dextrose 157.5 g

Agar 37.5g

Nipagen 50 mL (5g tegosept/50 mL 95% ethanol)

Instructions:

Fill a large pot with the starting water and put on full heat.

Fill the smaller pot with hydrating water.

Add sucrose and dextrose to the starting water and stir.

Add agar to the starting water when simmering and stir.

Put the corn flour and yeast in the hydrating water and stir.

Once the sugar/agar mix has turned golden and there are no longer clumps, add the corn flour/yeast mixture and stir.

Autoclave on liquid setting, sterilization time 20 min.

Keeping the food warm on the hot plate add the food additive or same volume/weight water for the control and stir.

Pour the food while liquid into individual vials.

Add nipagen to the surface of the food after 10 minutes when it has cooled significantly (to prevent mold on the food surface).

